# Spy Crickets: Use (or not) of heterospecific acoustic cues for anti-predator responses

**DOI:** 10.64898/2026.08.09.742598

**Authors:** Piper K. Zander, Ned A. Dochtermann

**Affiliations:** Department of Biological Sciences, North Dakota State University; Department of Biological Sciences, Cal Poly Humboldt

## Abstract

The ability of prey to eavesdrop on predator vocalizations is expected to increase survival by reducing detection and capture. Unfortunately, most research has been conducted in vertebrates, and little is known about this ability in invertebrates. We measured latency to emerge, overall activity, and shelter visits in wild-caught fall field crickets (*Gryllus pennsylvanicus*) in response to acoustic playback. Stimuli included multiple predator vocalizations, non-predator vocalizations, white noise, and a control. We predicted that crickets would reduce activity, spend more time in shelter, and freeze in response to stimuli representing greater risk. Contrary to our predictions, crickets traveled greater distances, spent more time moving, and spent less time in shelter in response to predator vocalizations versus controls. We did not, however, find clear differences in responses between predator vocalizations and other treatments. Our results suggest that crickets may not differentiate between the vocalizations of predators, non-predators, and other abrupt sounds. Consequently, eavesdropping may not be a viable method of assessing predation risk for this species and its general use remains unclear.

## Introduction

Acoustic signals are commonly used to communicate information between individuals, and can include information about resource availability, territory, mating, and predation risk (Dunlop et al., 2022). Although signals are typically directed towards conspecifics, they may be intercepted by members of other species, generally referred to as eavesdropping regardless of the modality of signals (Dunlop et al., 2022; Magrath et al., 2015). Eavesdropping on signals meant for another receiver is particularly common for alarm calls; i.e. signals that alert others to the presence of a predator and elicit anti-predator responses in their receivers (Magrath et al., 2015). Species have been observed to increase vigilance, flee, find shelter, and perform mobbing behaviors in response to heterospecific alarm calls (Lea et al., 2008; Rainey et al., 2004; Schmidt et al., 2008; Vitousek et al., 2007). Eavesdropping also occurs between prey and predators where prey species respond with anti-predator behaviors to the vocalizations of their predators (Blumstein et al., 2008; Deecke et al., 2002; Hettena et al., 2014; Rainey et al., 2004). The ability to eavesdrop on heterospecific acoustic signals, whether they are alarm calls or predator vocalizations, provides prey species with both immediate and long-term benefits: reduced detection, greater chance to avoid capture, reduced vigilance, and increased time spent foraging (Magrath et al., 2015).

Unfortunately, most research on prey eavesdropping on predator vocalizations focuses on vertebrate species (Hettena et al., 2014; Yack et al., 2020). This reflects the broader taxonomic bias present in animal behavior research (Dochtermann et al., 2026; Rosenthal et al., 2017). Studies that do investigate this process in invertebrates have mostly focused on flying arthropods (Dawson et al., 2004; Fournier et al., 2013; Jacobs et al., 2008; Kollross et al., 2023; Pollack, 2015; Ter Hofstede et al., 2010; Triblehorn et al., 2008; Yack et al., 2007, 2020). Flying Orthopterans, for example, have been observed to respond both behaviorally and physiologically to auditory cues associated with flying predators (Belovsky et al., 2011; Dawson et al., 2004; Schulze & Schul, 2001; ter Hofstede et al., 2010) and multiple species of flying crickets exhibit anti-predator behaviors when exposed to ultrasonic stimuli such as bat echolocation calls (Farris et al., 1998; Fukutomi & Ogawa, 2017; Fullard et al., 2005; Nolen & Hoy, 1986; Pollack, 2015; Römer & Holderied, 2020). However, many arthropods, including field crickets, spend much or all their time on the ground, and are preyed upon by both aerial and terrestrial predators (Adamo et al., 2013; Gawałek et al., 2014; Hedrick & Dill, 1993; Li et al., 2026; Niemelä et al., 2012). Little research has examined how ground-dwelling arthropods respond to predator vocalizations. Here, we investigated whether the fall field cricket, *Gryllus pennsylvanicus*, eavesdrops on heterospecific vocalizations, how they might use that information to assess predation risk, and how and if they adjust their antipredator behavior according to risk.

Crickets are consumed by many vertebrate predators including birds, rodents, and toads (Gawałek et al., 2014; Hedrick, 2000; Li et al., 2026; Niemelä et al., 2012). These predators often use acoustic signals to communicate with heterospecifics but do not vocalize before attacking their prey. Therefore, to successfully implement anti-predator behaviors that reduce their predation risk from such predators, crickets need to intercept acoustic signals made by predators prior to predation occurring. However, anti-predator behaviors are costly: increased vigilance decreases time spent foraging and reproducing and fleeing uses energy that must then be regained (Kats & Dill, 1998; Lima, 1998; Turbill & Stojanovski, 2018). To maximize time allocated to beneficial activities and reduce the chance of predation, prey are predicted to assess levels of predation risk (Helfman, 1989). Consistent with this prediction, crickets have been observed to alter their anti-predator behavior based on predation risk at both the individual and population level. For example, the wood cricket, *Nemobius sylvestris*, adjusts its anti-predator response to spider chemical cues based on perceived risk according to spider size and hunting strategy (Binz et al., 2014). Relatedly, western field cricket (*Gryllus integer*) populations under high risk of predation were found to be more cautious than crickets from populations with low predation risk when presented with a simulated predator cue (Kortet et al., 2007) and the European field cricket (*G. campestris*) responds to cues mimicking predator proximity (Li et al., 2026). Crickets may also be able to intercept acoustic signals produced by predators and interpret those signals to determine the level of risk they pose, using that information to adjust their behavior.

We investigated whether the fall field cricket, *G. pennsylvanicus*, can eavesdrop on heterospecific vocalizations and adjust anti-predator behavior in response to potential predation risk. We predicted that if crickets can assess predation risk from acoustic stimuli, then individuals should exhibit increased rates of freezing, reduced overall activity, and spend more time in shelter following the introduction of a predator cue. Conversely, crickets were predicted to not exhibit freezing, reductions in activity, or retreats to shelter following a non-predator cue.

## Methods

### Collection and Housing of Study Species

Adult male and female *G. pennsylvanicus* were collected in Fargo, North Dakota from August through October 2025. Upon capture crickets were housed individually in 0.6 L paper cups with cardboard egg carton for shelter and *ad libitum* water and food. Crickets were kept in a controlled environment with a reversed 12:12 h light/dark cycle at 27°C. As dealing with invertebrates, this research was not under review by an institutional review board but was conducted in accordance with ASAB/ABS guidelines.

### Behavioral Experiments

We conducted playback experiments on 30 wild-caught *G. pennsylvanicus* (F=22, M=8) to assess behavioral responses to predator acoustic cues, non-predator acoustic cues, white noise, and a control (see below). This skewed sex ratio was due to the outcome of opportunistic sampling. These experiments tested two components of behavior: emergence from shelter and anti-predator response. We used activity measurements and retreats to shelter as measures of anti-predator behavior. We selected measurements (Table 1) in accordance with prior studies that showed multiple cricket species demonstrate anti-predator behaviors including a cessation of calling, freezing, and fleeing in response to visual, auditory, and vibrational cues associated with predation (Adamo et al., 2013; Hedrick, 2013; Hedrick, 2000; Kortet et al., 2007). Additionally, crickets show changes in activity when exposed to chemicals produced by vertebrate and invertebrate predators (Binz et al., 2014; Bucklaew & Dochtermann, 2021; Dalos et al., 2022; Hoefler et al., 2012; Royauté & Dochtermann, 2017; Storm & Lima, 2008; Tanis et al., 2018). *G. pennsylvanicus*, for example, have demonstrated a reduction in movement and speed in the presence of chemical cues of a spider predator (Storm & Lima, 2008).

**Table 1.**
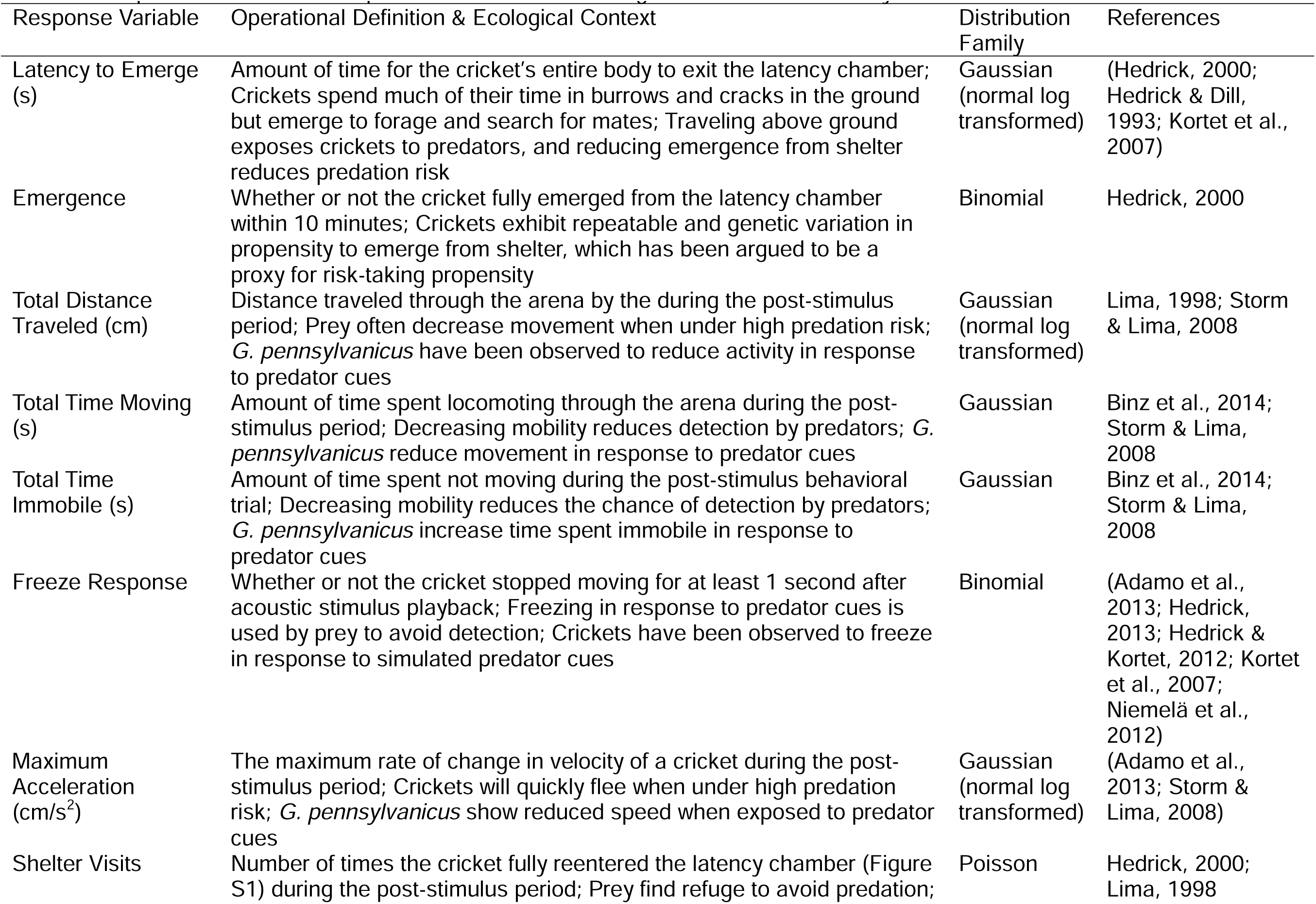
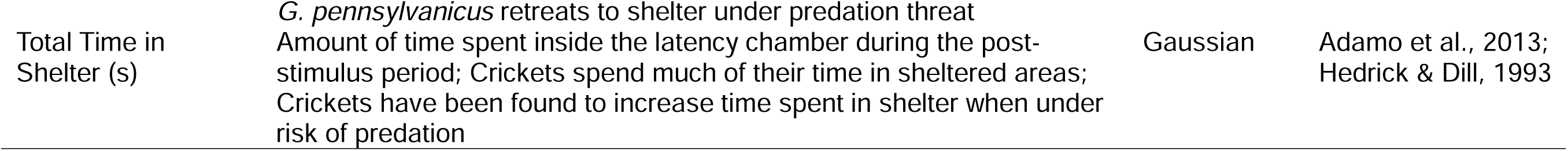
Response variables, their operational definitions, ecological relevance, and analysis families.

We conducted behavioral trials over a four-week period from late September to late October 2025. Testing was conducted in multiple groups based on capture date. Groups one and two consisted of nine crickets while group three consisted of 12 crickets. Each individual underwent a series of six trials wherein a cricket was presented with one of six acoustic stimuli during each trial (see below). To reduce the chance of habituation, trials were separated by at least 24 h. Stimulus order was determined via a six-by-six William’s square design to reduce the impact of carry-over effects between stimuli (Díaz-Uriarte, 2002; Dochtermann, 2010). Due to senescence and some crickets dying before completing all six trials, a total of 156 trials were conducted (Table 2).

**Table 2.**
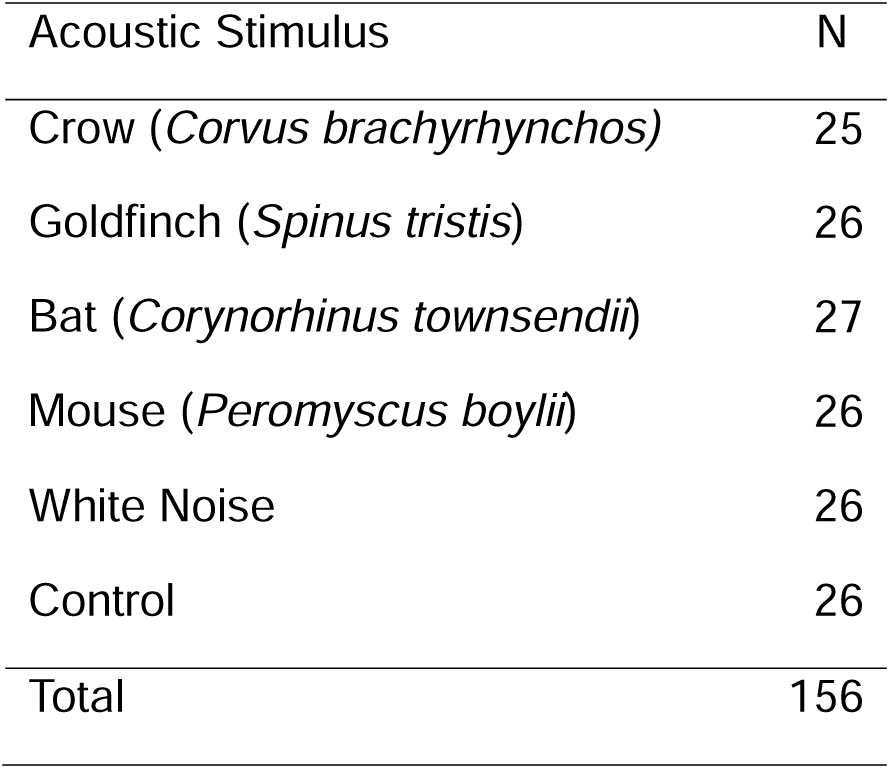
Number of individuals (N) tested for each stimulus type.

Prior to each trial, the focal cricket had its mass recorded and was placed into a small, 1.5 cm tall artificial burrow (∼20 cm^3^) with an opening towards one end (Figure S1). A sliding door was inserted through the sides of the opening to prevent the cricket from exiting before the start of the trial. The opening of the chamber was then slotted into the side of an opaque plastic arena (30 x 30 x 40 cm) with a camera (StreamCam, Logitech, Lausanne, Switzerland) and ultrasonic speaker (Avisoft Bioacoustics, Berlin, Germany) positioned 45.5 cm above the arena (Figure S2). Following a 90 s acclimation period, video recording was started and the sliding door was removed from the latency chamber. Once the cricket’s full body emerged from the chamber and the cricket was actively moving through the arena, an acoustic stimulus was played, and the cricket’s activity was recorded for the subsequent 5 minutes.

At the conclusion of the trial, or if the cricket did not fully emerge within 10 min, the cricket was returned to individual housing. The arena and latency chamber were sanitized with 70% ethanol between trials. Emergence and latency to emerge from shelter were directly recorded from trial videos (Table 1).

To assess responses of crickets to the acoustic stimuli, we processed each video recording using EthoVision XT (Noldus Information Technology, Wageningen, the Netherlands). We extracted the following measurements following video processing: total distance traveled, total time spent moving, total time spent immobile, latency to start moving after the stimulus, latency to stop moving after the stimulus, maximum acceleration, number of arena visits, and time spent in the arena (Table 2).

Number of arena visits was converted to number of shelter visits; time spent in the arena was converted into time spent in shelter (Table 1). We also measured the duration of freezing response by subtracting latency to stop moving after the stimulus from latency to start moving after the stimulus. These values were then converted into a binary freeze response (1 = froze; 0 = did not freeze) where freezing was reported as not moving for greater than one second after the stimulus was played. This is a conservative cutoff and well below the median duration of freezing for both males and females as previously reported (Hedrick & Kortet, 2012).

### Selection and Analysis of Acoustic Stimuli

Acoustic stimuli included predator and non-predator vocalizations in both ultrasonic (>20 kHz) and human-audible (<20 kHz) ranges. Crickets have tympanal ears located on the proximal tibia of each foreleg which can detect a broad range of frequencies, including those of the acoustic cues presented here (Yack, 2004). Female *G. pennsylvanicus* have the greatest phonotactic response to male calling songs between 4-6 kHz (Jeffery et al., 2005), corresponding with the average carrier frequency of male courtship and long-distance mating calls (∼4-5 kHz) (Harrison et al., 2013). The vocalizations of both predator and non-predator birds often overlap with these frequencies, suggesting that crickets can hear their calls. Additionally, male *G. pennsylvanicus* courtship calls contain high frequency ticks that extend into the ultrasonic range (Harrison et al., 2013). This, in addition to multiple other *Gryllus* species being capable of hearing ultrasonic bat calls (Farris et al., 1998; Fukutomi & Ogawa, 2017; Pollack & Martins, 2007), suggests that *G. pennsylvanicus* may also be able to hear and respond to ultrasonic predator vocalizations. Based on this known and potential auditory range, we examined the response to four potential predator vocalizations: two human-audible vocalizations and two ultrasonic vocalizations.

Human-audible stimuli: The acoustic stimuli included two bird calls: an insectivorous bird (American crow, *Corvus brachyrhynchos*) and a non-insectivorous bird (American goldfinch, *Spinus tristis*). Recordings were obtained from the Cornell Lab of Ornithology Macaulay Library (ML629047944, ML631185319, ML530790021, ML622920440, ML627261486, ML627872716). Crows readily consume arthropods, including ground-dwelling species, making them a likely predator of *G. pennsylvanicus* (Barrows & Schwarz, 1895; Johnson, 1994), and *G. pennsylvanicus* freeze for longer durations in response to playbacks of sounds produced by avian predators (calls and wingbeats) than to playbacks of sounds produced by non-avian predators (Brown, 2024). We predicted that if crickets can distinguish predation risk between avian species, they will exhibit increased anti-predator responses to calls of crows versus calls of goldfinches.

Ultrasonic vocalizations: We used the search-phase echolocation calls of Townsend’s big-eared bats (*Corynorhinus townsendii*) and social vocalizations of brush mice (*Peromyscus boylii*, Petric & Kalcounis-Rueppell, 2013). Townsend’s big-eared bats capture insects from the air and glean insects from foliage, but have not been found to consume *Gryllus* species, making them a lower risk predator for *G. pennsylvanicus* (Edens et al., 2023). Brush mice are known to consume arthropods, including crickets, and have an overlapping range with *G. pennsylvanicus* (Kalcounis-Rueppell & Spoon, 2009; Weissman et al., 1980), although they do not co-occur in North Dakota. *G. pennsylvanicus* in North Dakota may possess the ability to respond to *P. boylii* due to their ecological overlap with other *Peromyscus* species and because many prey species can respond to vocalizations of predators with which they share an evolutionary history despite a lack of current exposure (Hettena et al., 2014).

For each of the four species, three representative samples were used. Two seconds of vocalizations were isolated from each bat and crow recording, and 0.5 seconds of vocalizations were isolated from each mouse and goldfinch recording. This discrepancy in recording duration was due to variation in call length between species; bat and crow calls were longer than mouse and goldfinch calls. Low frequency background noise was reduced from each recording using the “Noise Reduction” feature in the software Audacity (version 3.7.5; https://www.audacityteam.org/). In addition to the vocalization playbacks, three two-second-long clips of white noise were generated using Avisoft-SASLab Pro software (version 5.3.1; Avisoft Bioacoustics, Berlin, Germany). Three two-second-long recordings were also taken in the trial room for use as a control treatment.

Acoustic stimuli playback was calibrated and analyzed for spectral and amplitude characteristics prior to the behavioral trials. An ultrasonic speaker (Avisoft Bioacoustics, Berlin, Germany) was placed above the arena at the same height as it would be during the behavioral trials (Figure S2). A sound level meter (PSPL25, Pyle USA, Brooklyn, NY, USA) and ultrasonic microphone (CM16/CMPA, Avisoft Bioacoustics, Berlin, Germany) were placed in the center of the bottom of the arena to measure loudness and acoustic characteristics of playbacks at a position approximating that of where a cricket would be during trials. The microphone was connected to an Avisoft-UltraSoundGate 416H A/D converter; the speaker was connected to an Avisoft-UltraSoundGate Player 116 D/A converter (Avisoft Bioacoustics, Berlin, Germany). Each audio clip was played from, and recorded onto, a laptop computer using Avisoft RECORDER USGH software (version 4.4.1; Avisoft Bioacoustics, Berlin, Germany): ultrasonic stimuli were recorded at a 16-bit sampling frequency of 250 kHz, human-audible stimuli were recorded at a 16-bit sampling frequency of 48 kHz. We measured spectral and amplitude characteristics including peak amplitude, best frequency (peak frequency at peak amplitude), and bandwidth at 10 dB below the peak frequency. The volume of each audio clip was adjusted to 80-90 dB given background noise and the approximate distance from the subject to the speaker following Fischer et al. (2013).

### Statistical Analysis

To test for differences in cricket responses, we built linear mixed effects models (LMMs) and generalized linear mixed effects models (GLMMs) for each response variable (Table 2) using the lme4 package (version 1.1.37; Bates et al., 2015) in R (version 4.5.0; R Core Team, 2025). Emergence time, total distance traveled, total time spent moving, total time spent immobile, maximum acceleration, and total time spent in shelter were analyzed as having Gaussian distributed residuals (Table 1). Emergence and freeze response were analyzed as being Binomial distributed; shelter visit count was analyzed as being Poisson distributed (Table 1). Treatment was included as a fixed effect in all models except for where emergence or emergence time were the response variables (these behavioral responses occurred prior to treatment application).

We initially planned to include trial number, sex, mass, number of days in captivity, injury status, temperature, time of trial, sequence ID, and batch number as fixed effects. However, due to our final sample size and resulting statistical separation, we fit reduced models including only sex, mass, days in captivity, injury status, temperature and treatment. The reduced models were fit to untransformed data for emergence time, total distance traveled, total time spent moving, total time spent immobile, maximum acceleration, and total time spent in shelter and Q-Q plots visibly examined. Although linear mixed-effects models are generally robust to assumption violation (Schielzeth et al., 2020), when substantial deviation of residuals from expectations was identified in Q-Q plots, we reran models on log transformed data (Table 1). Trial number was also included as a fixed effect for models of emergence and emergence time. Individual ID was included as a random effect in all models. Mass, temperature, trial number, and days in captivity were centered for analyses.

Significance of effects was determined using the lmerTest package (version 3.1.3; Kuznetsova et al., 2017). We also performed specific pre-planned contrasts between treatments using single-step p-value adjustment using the multcomp package (version 1.4.30; Hothorn et al., 2008).

Finally, because we had up to six repeated measures per individual, we calculated unadjusted repeatabilities (Nakagawa et al., 2026) using the rptR package (version 0.9.23; Stoffel et al., 2017). We estimated the uncertainties of these repeatabilities via bootstrapping (n = 1000).

## Results

### Pre-stimulus Behavioral Measures

Crickets emerged from shelter in 104 of 156 behavioral trials. Trial number, mass, days in captivity, injury status, and temperature did not have a significant effect on emergence from shelter or latency to emerge from shelter (Table S1, S3). Additionally, sex did not have a significant effect on emergence (Table S3). In contrast, sex significantly affected emergence time (F_df= 1, 24.312_ = 4.3077, p = 0.049; Figure 1, Table S1). Male crickets took significantly, albeit only very slightly, longer to emerge from shelter compared to female crickets (difference = 1.1 seconds, t = 2.075, p = 0.049; Figure 1, Table S2).

**Figure 1.**
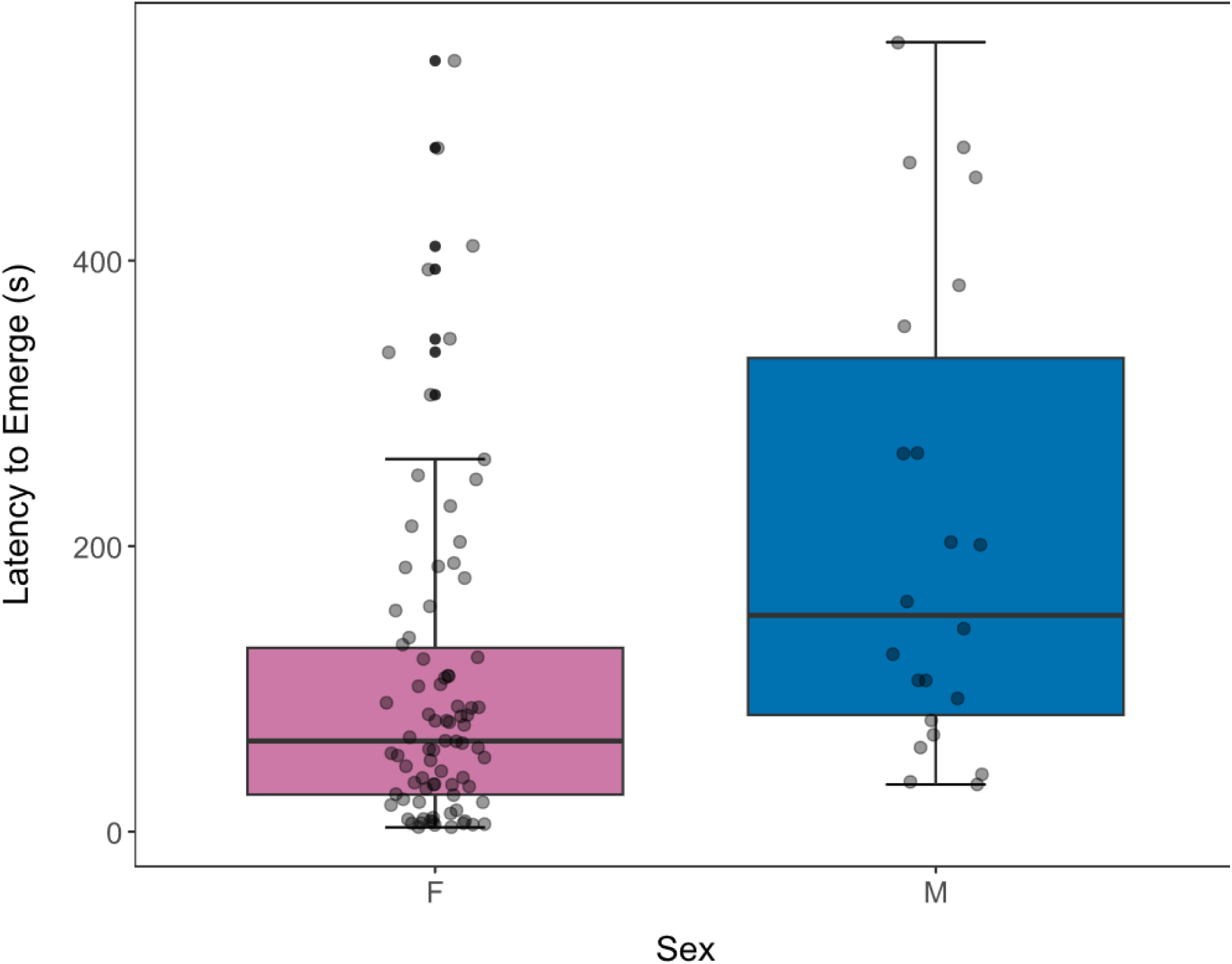
Boxplots of latencies to emerge in seconds for female (red) and male (blue) crickets. The central line represents the median and the lower and upper bars represent 1^st^ and 3^rd^ quartiles. The error bars show the range.

### Post-stimulus Behavioral Measures

Across the 104 trials in which crickets emerged from shelter, treatment, sex, mass, injury status, days in captivity, and temperature did not result in differences in total time spent immobile, maximum acceleration, or number of shelter visits (Table S1, S3). Treatment, sex, mass, days in captivity, and temperature did not affect freeze response (Table S3). Injury status affected freeze response, with crickets that were missing one or both jumping legs being more likely to freeze in response to acoustic stimuli (z = 2.411, p = 0.016; Table S3). Additionally, sex, mass, injury status, days in captivity, and temperature had no effect on total distance traveled, total time spent moving, or time spent in shelter (Table S1).

We found that treatment had a significant effect on total distance traveled (F_df= 5, 75.167_ = 2.610, p = 0.031; Figure 2a, Table S1) with total distance traveled after playback of bat echolocation (difference = 0.7 cm, z = -2.779, p = 0.048; Figure 2a, Table S4), mouse calls (difference = 0.7 cm, z = -3.058, p = 0.021; Figure 2a, Table S4), and white noise (difference = 0.7 cm, z = -2.968, p = 0.028; Figure 2a, Table S4) being greater than after control stimuli. We also found that treatment had a significant effect on total time spent moving (F_df= 5, 75.861_ = 2.763, p = 0.024; Figure 2b, Table S1) with total time spent moving after playback of mouse calls being greater than after control stimuli (difference = 53.5 seconds, z = -2.853, p = 0.039; Figure 2b, Table S4). Total time spent moving after playback of white noise was also greater than after control stimuli (difference = 56.4 seconds, z = -3.138, p = 0.016; Figure 2b, Table S4).

**Figure 2.**
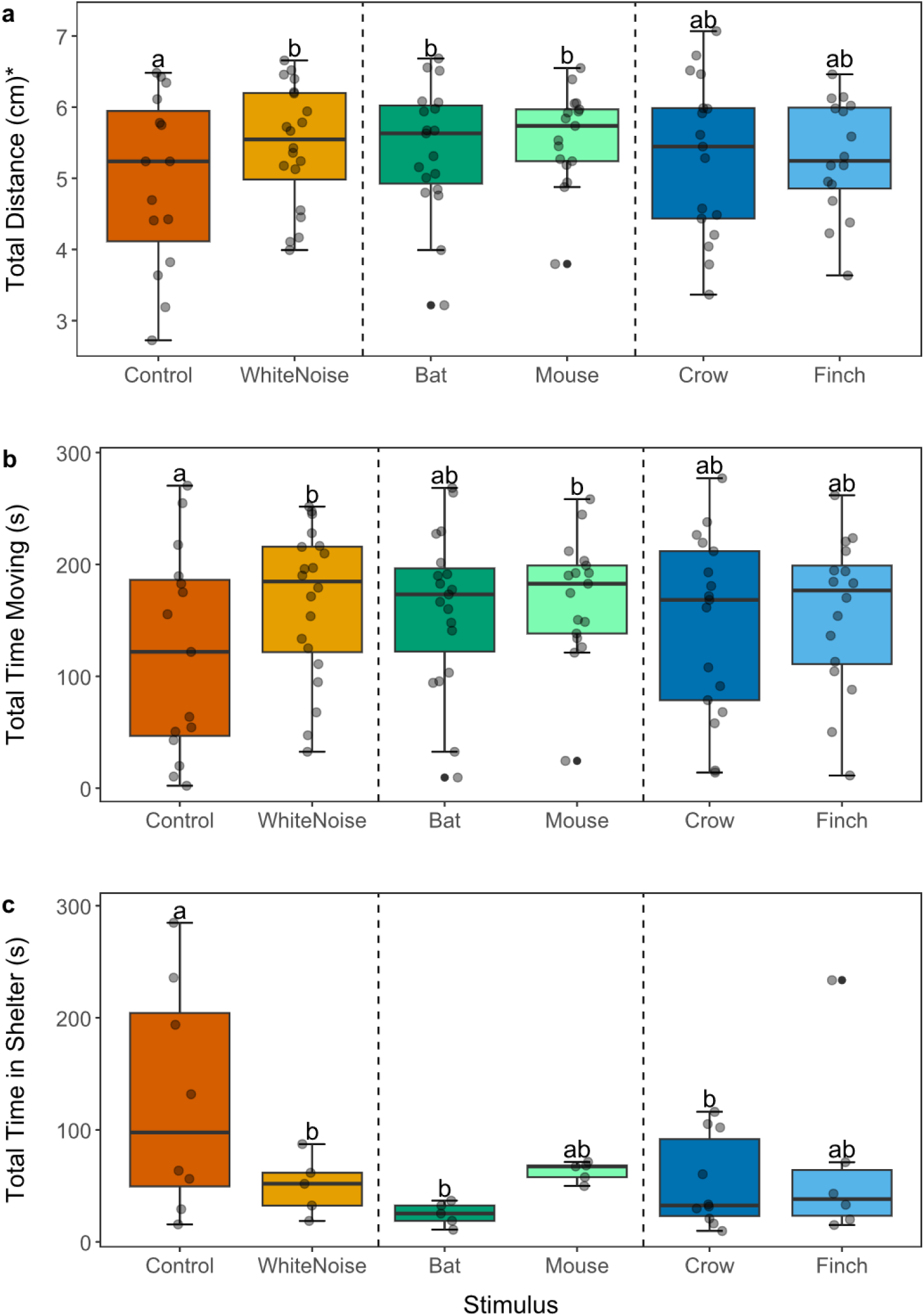
Boxplots of a) total distance traveled (log transformed), b) total time moving, and c) total time spent in shelter for crickets across stimuli. The central line represents the median and the lower and upper bars represent 1^st^ and 3^rd^ quartiles. The error bars show the range. Bars with different letters above them are significantly different from one another within panels (p < 0.05). *indicates log transformed values.

Time spent in shelter was also significantly affected by treatment (F_df= 5, 19.299_ = 3.924, p = 0.013; Figure 2c, Table S1). Specifically, crickets spent significantly less time in shelter after playback of bat echolocation (difference = 127.5 seconds, z = 3.966. p <0.001; Figure 2c, Table S4), crow calls (difference = 84.2 seconds, z = 3.133, p = 0.016; Figure 2c, Table S4), and white noise (difference = 90.2 seconds, z = 2.878, p = 0.036; Figure 2c, Table S4) when compared to the control. We found no statistically significant differences in response between non-control treatments (Table S4).

### Behavioral Repeatability

Repeatability ranged from low (number of shelter visits, τ = 0.197) to well above the average for behaviors (emergence, τ = 0.614; Figure 3, Table S5). Fixed effects explained little of the observed variation in any of the behaviors (Table S5). Full variance component estimates and associated uncertainties are reported in Table S5.

**Figure 3.**
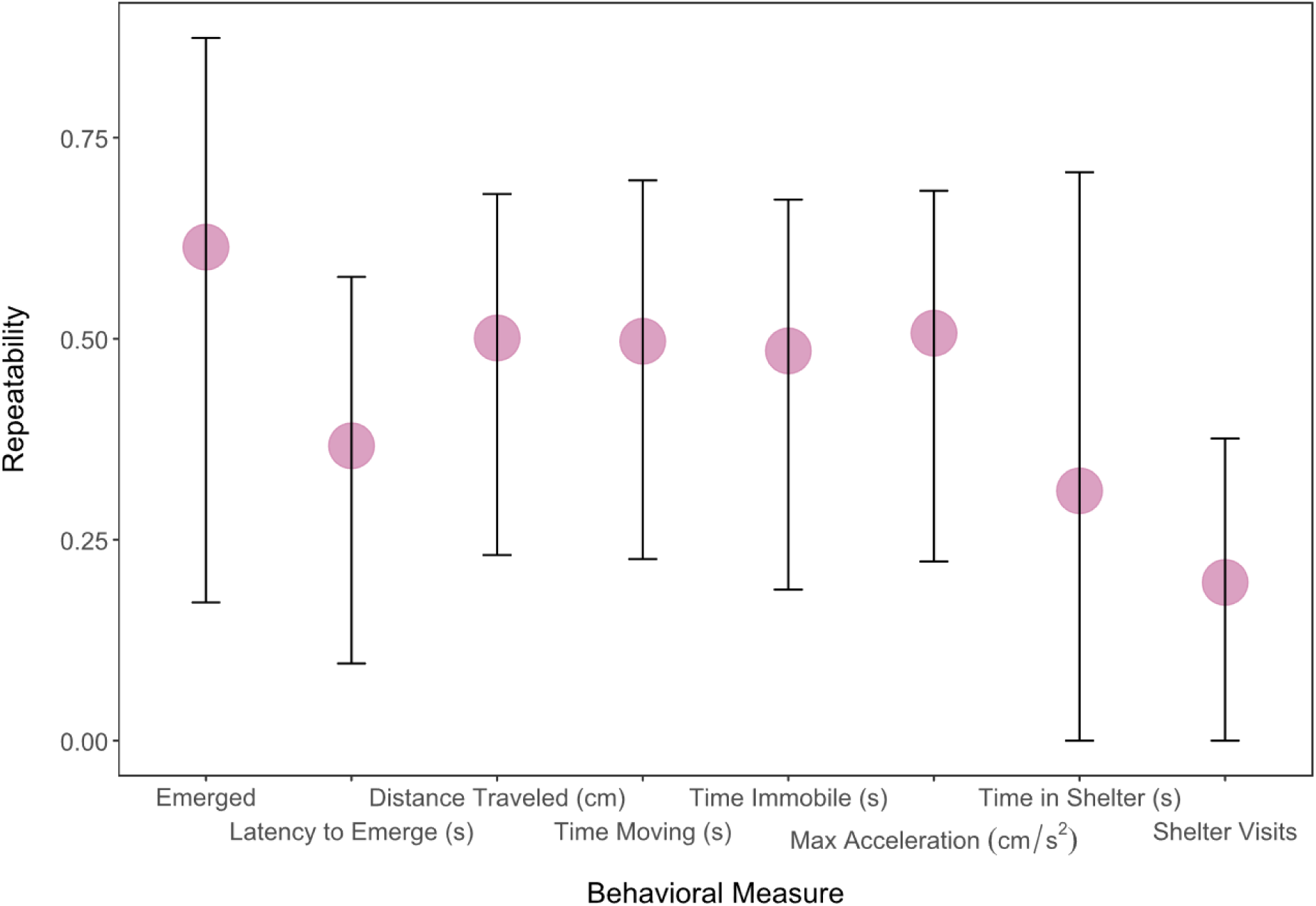
Adjusted repeatability values for each behavioral measure. Error bars show the upper and lower 95% confidence intervals.

## Discussion

### Pre-stimulus Behavioral Measures

Behavioral measures taken prior to the introduction of acoustic stimuli were not significantly affected by trial number, mass, days in captivity, injury status, or temperature. We did, however, find a slight significant effect of sex on latency to emerge from shelter with male crickets taking longer to emerge compared to female crickets.

Surprisingly, these findings contradict previous studies that found no difference in emergence time between male and female crickets (A. V. Hedrick & Kortet, 2012; Wilson et al., 2010). This discrepancy may be due to our study having a small sample size of males (n = 8) compared to females (n = 22). Alternatively, the slight difference in emergence time may not be biologically significant despite its statistical significance. We did not find an effect of sex on whether an individual emerged from shelter.

### Post-stimulus Behavioral Measures

Our results demonstrate that crickets respond to acoustic stimuli, but not as expected. We predicted that crickets would reduce activity, exhibit a longer freezing response, and retreat to shelter following playback of predator vocalizations. Contrary to our predictions, we found no effect of treatment on time spent immobile, maximum acceleration, freeze response, or number of shelter visits. We did find that freeze response was affected by injury status. Crickets that were missing one or both jumping legs were more likely to freeze in response to acoustic stimuli, likely due to the reduced mobility of these crickets.

We found that treatment, versus control stimuli, did have an effect on total distance traveled, time spent moving, and time spent in shelter. Crickets traveled significantly greater distances after exposure to bat and mouse calls compared to control recordings. Time spent moving significantly increased in response to mouse calls. We also found that crickets exposed to bat or crow calls spent significantly less time in shelter compared to the control. Additionally, we found that crickets exposed to white noise spent significantly more time moving and significantly less time in shelter when compared to the control. The non-predator treatment of goldfinch calls did not have a significant effect on any response variables. We found no discernable difference in response between predator vocalizations, white noise, or non-predator vocalizations. Crickets exhibited above-average repeatability in latency to emerge and below-average repeatability in number of shelter visits. This indicates that individual crickets are relatively consistent in how long they take to emerge from shelter and are less consistent in the number of times they return to shelter.

Previous studies on antipredator behavior in crickets, including *Gryllus* species, have shown responses of freezing, fleeing, finding and remaining in shelter, and reducing activity following the presentation of predator cues (Adamo et al., 2013; Binz et al., 2014; Hedrick, 2013; Hoefler et al., 2012; Kortet et al., 2007; Storm & Lima, 2008; Tanis et al., 2018). *G. texensis* have been observed to spend more time in shelter when exposed to a mock predator compared to non-exposed crickets and flee in the presence of a live predator (Adamo et al., 2013). In *G. pennsylvanicus,* exposure to chemical predator cues results in a significant increase in time spent immobile and reduction in movement speed (Storm & Lima, 2008). Brown (2024) found a non-significant effect of avian predator playbacks on *G. pennsylvanicus* freezing response where crickets exposed to avian cues froze for longer than crickets exposed to the cues of other predators. We did not, however, find a similar response toward predator vocalizations. Surprisingly, we instead found that crickets showed increased activity and spent less time in shelter in response to predator vocalizations. This is consistent with findings in other Gryllid crickets that exposure to chemical cues of vertebrate predators with active hunting strategies elicits an increase in activity (Bucklaew & Dochtermann, 2021; Royauté & Dochtermann, 2017; Royauté et al., 2019). We also found no effect of predator vocalizations on the occurrence of a freeze response.

These discrepancies between our findings and previous studies suggests that *G. pennsylvanicus* may not recognize predator vocalizations as indicators of predation risk. In support of this suggestion, crickets did not show a significantly different response to goldfinch calls compared to other stimuli. As goldfinches are non-insectivorous, we predicted that playback of their calls would not elicit anti-predator behaviors, and that response to their calls would differ from response to predator vocalizations. As we did not find a difference in response to predator and non-predator vocalizations, our findings suggest that as well as not perceiving risk from predator vocalizations, crickets may not differentiate between vocalizations of other species in general. We also found, however, that response to white noise was similar to that of predator cues. This similarity in response may suggest that crickets do not distinguish between heterospecific vocalizations and other loud, novel sounds.

Eavesdropping on predator vocalizations is observed in many species. For example, harbor seals differentiate between the calls of predatory transient and non-predatory resident orcas (Deecke et al., 2002), yellow-bellied marmots increase vigilance in response to avian and terrestrial predator calls (Blumstein et al., 2008), and for many flying insects, bat echolocation induces negative phonotaxis (Pollack, 2015). The ability to distinguish between the sounds of predators and non-predators confers the benefit of reducing predation risk for prey, as it allows them to implement anti-predator responses before being detected by a predator. Despite these benefits, our results do not support field crickets eavesdropping on predator vocalizations. This may be due to the energetic cost of anti-antipredator behavior. Vigilance reduces time available for other beneficial activities such as mating and foraging, therefore, responding to every social call a predator makes with increased vigilance may negatively impact overall growth and reproduction (Lima, 1998). Here, we did not find a reduction in activity associated with increased vigilance in crickets in response to predator calls, suggesting that the energetic cost of eavesdropping and responding may outweigh the benefits in reducing predation risk. Relatedly, we did not find an increased freezing response following exposure to white noise. This lack of startle response is surprising, as white noise is a novel sound for these crickets. However, the lack of response may be indicative of a trade-off where responding to every novel sound with anti-predator behavior results in a reduction in time available for other activities (Magrath et al., 2015). Instead, crickets may prioritize other beneficial activities over vigilance.

Our results suggest that eavesdropping on predator vocalizations may not be a viable method of assessing predation risk for *G. pennsylvanicus*. This may be due to the energetic cost associated with remaining vigilant, or it may be due to how predators hunt. Most predators do not vocalize while they are actively hunting, and if hunting predators are usually silent, prey may not recognize their vocalizations as indicative of an immediate risk (Schmidt et al., 2008). Therefore, additional cues may be needed to evoke a behavioral response in crickets. A study on desert locusts (*Schistocerca gregaria*) found that time in shelter and stress response significantly increased upon exposure to live birds and bird call playback, but not to playback of bird calls only (Kollross et al., 2023). The presence of a live predator, or visual predator cues, may be needed to induce anti-predator behavior in *G. pennsylvanicus*. Additionally, changes in air movement and vibrational and chemical cues may be more relevant for indicating predator presence than vocalizations. Cricket cerci are covered in hairs that are sensitive to air currents, and changes in air movement associated with an approaching predator can trigger escape responses (Fukutomi & Ogawa, 2017; Kiuchi et al., 2023; Lagos, 2017). The movement of an approaching terrestrial predator also results in substrate-borne vibrations which have been shown to elicit anti-predator responses in field crickets (Hedrick, 2000; Kiuchi et al., 2023; Kortet et al., 2007; Li et al., 2026). Additionally, predators produce chemical signals that crickets are able to sense and respond to (Binz et al., 2014; Bucklaew & Dochtermann, 2021; Dalos et al., 2022; Hoefler et al., 2012; Pears et al., 2018; Royauté & Dochtermann, 2017; Storm & Lima, 2008; Tanis et al., 2018). Future research should explore how *G. pennsylvanicus* respond to non-acoustic cues associated with varying levels of predation threat.

Elsewhere, prey species have been observed to eavesdrop on the vocalizations of predators, even when those predators do not vocalize immediately prior to attacking (Blumstein et al., 2008; Deecke et al., 2002; Hettena et al., 2014). This ability to recognize predation risk from heterospecific vocalizations confers fitness benefits via increasing survivability. Surprisingly, we did not find evidence to support *G. pennsylvanicus* utilizing eavesdropping as a method of assessing predation risk. These results suggest that eavesdropping on heterospecific vocalizations may not be widespread across taxa, and future studies should further investigate the use of eavesdropping on acoustic cues in invertebrates.

## Data Availability

All data and analysis code are available at: link.

## Acknowledgements

We thank the NSF CHANGE RaMP program for supporting this project (DBI-2216605). Thank you to Dr. Erin Gillam, Dr. Matina Kalcounis-Rueppell, and the Cornell Lab of Ornithology | Macaulay Library for supplying acoustic playback materials. We also thank L. Frias for extensive, helpful discussions regarding question and method development.

