## Supplemental Materials for "Spy Crickets: Use (or not) of heterospecific acoustic cues for anti-predator responses"

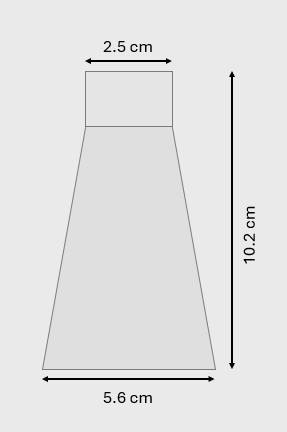

Figure S1. Top-down schematic of latency chamber.

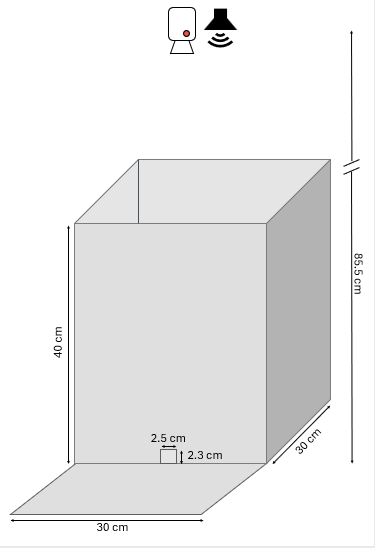

Figure S2. Schematic for behavioral trial set-up including the arena, camera, and speaker.

| Table S1. Summary of ANOVA results for latency to emerge, total distance traveled, total time moving, total time immobile, total time in shelter, and maximum acceleration. Bold values indicate statistically significant effects (p<0.05). * indicates log transformed variables. | | | | | | | |
| --- | --- | --- | --- | --- | --- | --- | --- |
| Response Variable |  | Sum Sq | Mean Sq | NumDF | DenDF | F-value | p-value |
| Latency to Emerge* | Mass | 0.0786 | 0.0786 | 1 | 33.494 | 0.0859 | 0.7713 |
|  | Temperature | 2.2051 | 2.2051 | 1 | 89.206 | 2.4088 | 0.12419 |
|  | Days in Captivity | 3.6902 | 3.6902 | 1 | 21.033 | 4.0311 | 0.05769 |
|  | Trial Number | 0.2518 | 0.2518 | 1 | 94.153 | 0.275 | 0.60121 |
|  | Injury Status | 0.933 | 0.933 | 1 | 23.363 | 1.0192 | 0.32305 |
|  | Sex | 3.9434 | 3.9434 | 1 | 24.312 | 4.3077 | **0.04868** |
| Total Distance Traveled* | Mass | 0.0509 | 0.05091 | 1 | 42.31 | 0.1175 | 0.73351 |
|  | Temperature | 0.0156 | 0.01564 | 1 | 86.703 | 0.0361 | 0.84978 |
|  | Days in Captivity | 0.1579 | 0.15789 | 1 | 38.961 | 0.3643 | 0.54964 |
|  | Injury Status | 1.5137 | 1.51367 | 1 | 27.32 | 3.4924 | 0.07241 |
|  | Sex | 0.3119 | 0.31187 | 1 | 25.892 | 0.7196 | 0.40406 |
|  | Treatment | 5.6563 | 1.13126 | 5 | 75.167 | 2.6101 | **0.03129** |
| Total Time Moving | Mass | 166 | 166.2 | 1 | 43.269 | 0.0641 | 0.80134 |
|  | Temperature | 35 | 34.7 | 1 | 87.044 | 0.0134 | 0.9082 |
|  | Days in Captivity | 96 | 96.1 | 1 | 39.723 | 0.0371 | 0.84832 |
|  | Injury Status | 6695 | 6695.3 | 1 | 28.009 | 2.5817 | 0.11932 |
|  | Sex | 2118 | 2118.1 | 1 | 26.77 | 0.8167 | 0.3742 |
|  | Treatment | 35831 | 7166.2 | 5 | 75.861 | 2.7633 | **0.02396** |
| Total Time Immobile | Mass | 14.3 | 14.3 | 1 | 41.03 | 0.0048 | 0.94497 |
|  | Temperature | 11673 | 11673.4 | 1 | 86.748 | 3.9329 | 0.05051 |
|  | Days in Captivity | 709.7 | 709.7 | 1 | 37.101 | 0.2391 | 0.62772 |
|  | Injury Status | 12175 | 12174.7 | 1 | 25.875 | 4.1018 | 0.05327 |
|  | Sex | 857.6 | 857.6 | 1 | 25.181 | 0.2889 | 0.59562 |
|  | Treatment | 9011.7 | 1802.3 | 5 | 74.727 | 0.6072 | 0.69456 |
| Total Time in Shelter | Mass | 2156 | 2156.5 | 1 | 12.609 | 0.8215 | 0.38175 |
|  | Temperature | 4693 | 4693 | 1 | 26.049 | 1.7877 | 0.19277 |
|  | Days in Captivity | 4399 | 4399.3 | 1 | 13.787 | 1.6758 | 0.21675 |
|  | Injury Status | 4659 | 4659.2 | 1 | 10.241 | 1.7748 | 0.21167 |
|  | Sex | 8470 | 8469.6 | 1 | 11.295 | 3.2263 | 0.09923 |
|  | Treatment | 51509 | 10301.8 | 5 | 19.299 | 3.9242 | **0.01273** |
| Maximum Acceleration* | Mass | 0.0313 | 0.03127 | 1 | 44.538 | 0.3203 | 0.57427 |
|  | Temperature | 0.0153 | 0.0153 | 1 | 87.125 | 0.1567 | 0.69316 |
|  | Days in Captivity | 0.0005 | 0.0005 | 1 | 41.381 | 0.0051 | 0.94327 |
|  | Injury Status | 0.3898 | 0.38976 | 1 | 29.362 | 3.9924 | 0.05504 |
|  | Sex | 0.0185 | 0.01845 | 1 | 27.609 | 0.189 | 0.66715 |
|  | Treatment | 0.6654 | 0.13309 | 5 | 76.39 | 1.3633 | 0.24755 |

| Table S2. Summary of linear mixed-effects model results for latency to emerge, total distance traveled, total time moving, total time immobile, total time in shelter, and maximum acceleration. Bold values indicate statistically significant effects (p<0.05). * indicates log transformed variables. | | | | | | |
| --- | --- | --- | --- | --- | --- | --- |
| Response Variable |  | Estimate | SE | DF | t-value | p-value |
| Latency to Emerge* | Intercept | 4.23609 | 0.23962 | 18.41917 | 17.679 | **5.26E-13** |
|  | Mass | 0.40001 | 1.36501 | 33.4941 | 0.293 | 0.7713 |
|  | Temp | -0.44825 | 0.28882 | 89.20641 | -1.552 | 0.1242 |
|  | In Captivity | 0.04052 | 0.02018 | 21.03265 | 2.008 | 0.0577 |
|  | Test Number | 0.03708 | 0.0707 | 94.1532 | 0.524 | 0.6012 |
|  | Injured Status (yes) | -0.43011 | 0.42603 | 23.36264 | -1.01 | 0.323 |
|  | Sex (M) | 1.11062 | 0.53511 | 24.31196 | 2.075 | **0.0487** |
| Total Distance Traveled* | Intercept | 4.95809 | 0.25594 | 45.5871 | 19.372 | **< 2e-16** |
|  | Mass | -0.36447 | 1.06347 | 42.31025 | -0.343 | 0.73351 |
|  | Temperature | -0.03862 | 0.20331 | 86.70293 | -0.19 | 0.84978 |
|  | Days in Captivity | -0.00897 | 0.01487 | 38.96127 | -0.604 | 0.54964 |
|  | Injury Status (yes) | -0.65493 | 0.35046 | 27.31958 | -1.869 | 0.07241 |
|  | Sex (M) | -0.37205 | 0.4386 | 25.89223 | -0.848 | 0.40406 |
|  | Bat | 0.66731 | 0.24009 | 75.45965 | 2.779 | **0.00687** |
|  | Crow | 0.36632 | 0.23749 | 73.46965 | 1.542 | 0.12725 |
|  | Finch | 0.54364 | 0.25058 | 75.94688 | 2.17 | **0.03317** |
|  | Mouse | 0.74173 | 0.24255 | 74.77483 | 3.058 | **0.00309** |
|  | White Noise | 0.69017 | 0.23253 | 73.13618 | 2.968 | **0.00405** |
| Total Time Moving | Intercept | 125.512 | 19.7087 | 46.9573 | 6.368 | **7.47E-08** |
|  | Mass | -20.7282 | 81.8756 | 43.2688 | -0.253 | 0.80134 |
|  | Temperature | -1.8173 | 15.7143 | 87.0443 | -0.116 | 0.9082 |
|  | Days in Captivity | -0.2203 | 1.1443 | 39.7226 | -0.193 | 0.84832 |
|  | Injury Status (yes) | -43.2957 | 26.9458 | 28.0093 | -1.607 | 0.11932 |
|  | Sex (M) | -30.472 | 33.7177 | 26.7701 | -0.904 | 0.3742 |
|  | Bat | 50.3856 | 18.5685 | 76.137 | 2.713 | **0.00823** |
|  | Crow | 22.886 | 18.3685 | 74.2075 | 1.246 | 0.21671 |
|  | Finch | 45.6171 | 19.3789 | 76.6326 | 2.354 | **0.02114** |
|  | Mouse | 53.5232 | 18.7594 | 75.4867 | 2.853 | **0.00558** |
|  | White Noise | 56.4377 | 17.9857 | 73.8792 | 3.138 | **0.00244** |
| Total Time Immobile | Intercept | 102.202 | 20.8569 | 45.7031 | 4.9 | **1.24E-05** |
|  | Mass | -6.0137 | 86.5881 | 41.0301 | -0.069 | 0.945 |
|  | Temperature | 33.2742 | 16.7785 | 86.7484 | 1.983 | 0.0505 |
|  | Days in Captivity | 0.5914 | 1.2094 | 37.1005 | 0.489 | 0.6277 |
|  | Injury Status (yes) | 57.5247 | 28.4033 | 25.8746 | 2.025 | 0.0533 |
|  | Sex (M) | 19.0994 | 35.5315 | 25.1806 | 0.538 | 0.5956 |
|  | Bat | 17.1552 | 19.8559 | 74.997 | 0.864 | 0.3904 |
|  | Crow | 19.5269 | 19.6467 | 72.9785 | 0.994 | 0.3236 |
|  | Finch | 3.3439 | 20.7212 | 75.578 | 0.161 | 0.8722 |
|  | Mouse | -7.0577 | 20.0616 | 74.3504 | -0.352 | 0.726 |
|  | White Noise | -0.3615 | 19.238 | 72.6238 | -0.019 | 0.9851 |
| Total Time in Shelter | Intercept | 147.053 | 24.459 | 26.764 | 6.012 | **2.12E-06** |
|  | Mass | 136.83 | 150.969 | 12.609 | 0.906 | 0.38175 |
|  | Temperature | -32.573 | 24.362 | 26.049 | -1.337 | 0.19277 |
|  | Days in Captivity | 1.972 | 1.523 | 13.787 | 1.295 | 0.21675 |
|  | Injury Status (yes) | -49.03 | 36.803 | 10.241 | -1.332 | 0.21167 |
|  | Sex (M) | 93.571 | 52.094 | 11.295 | 1.796 | 0.09923 |
|  | Bat | -127.54 | 32.161 | 17.728 | -3.966 | **0.00093** |
|  | Crow | -84.249 | 26.887 | 21.137 | -3.133 | **0.00499** |
|  | Finch | -83.261 | 31.705 | 21.756 | -2.626 | **0.01551** |
|  | Mouse | -80.292 | 31.601 | 16.526 | -2.541 | **0.02143** |
|  | White Noise | -90.162 | 31.322 | 16.95 | -2.878 | **0.01045** |
| Maximum Accleration* | Intercept | 2.41575 | 0.12214 | 47.33194 | 19.779 | **<2e-16** |
|  | Mass | 0.2873 | 0.50764 | 44.53795 | 0.566 | 0.5743 |
|  | Temperature | -0.03824 | 0.09659 | 87.12521 | -0.396 | 0.6932 |
|  | Days in Captivity | 0.00051 | 0.0071 | 41.38131 | 0.072 | 0.9433 |
|  | Injury Status (yes) | -0.33479 | 0.16755 | 29.36202 | -1.998 | 0.055 |
|  | Sex (M) | -0.09118 | 0.20975 | 27.60854 | -0.435 | 0.6671 |
|  | Bat | 0.1554 | 0.11397 | 76.67642 | 1.363 | 0.1767 |
|  | Crow | 0.03572 | 0.11272 | 74.78762 | 0.317 | 0.7522 |
|  | Finch | 0.12753 | 0.11895 | 77.10938 | 1.072 | 0.287 |
|  | Mouse | 0.2239 | 0.11514 | 76.01134 | 1.945 | 0.0555 |
|  | White Noise | 0.22576 | 0.11037 | 74.47592 | 2.046 | **0.0443** |

| Table S3. Summary of generalized linear mixed-effect model results for emergence, freeze response, and shelter visits. Bold values indicate statistically significant effects (p<0.05). * indicates log transformed variables. | | | | | |
| --- | --- | --- | --- | --- | --- |
| Response Variable |  | Estimate | SE | z-value | p-value |
| Emergence (binary) | Intercept | 2.92893 | 1.08447 | 2.701 | **0.00692** |
|  | Mass | -5.78237 | 4.35151 | -1.329 | 0.18391 |
|  | Temp | 0.44666 | 0.7704 | 0.58 | 0.56206 |
|  | In Captivity | -0.06059 | 0.06899 | -0.878 | 0.3798 |
|  | Injured Status | -1.0715 | 1.34648 | -0.796 | 0.42616 |
|  | Sex (M) | -3.63324 | 1.94751 | -1.866 | 0.0621 |
|  | Trial Number | 0.34864 | 0.2111 | 1.652 | 0.09864 |
| Freeze Response (binary) | Intercept | -2.1441 | 0.791723 | -2.708 | **0.00677** |
|  | Mass | 0.099299 | 2.501101 | 0.04 | 0.96833 |
|  | Temp | 0.978873 | 0.877304 | 1.116 | 0.26452 |
|  | In Captivity | -0.00432 | 0.032937 | -0.131 | 0.89562 |
|  | Injured Status | 1.536523 | 0.637265 | 2.411 | **0.0159** |
|  | Sex (M) | 0.894899 | 0.977723 | 0.915 | 0.36004 |
|  | Bat | 0.06944 | 0.979531 | 0.071 | 0.94348 |
|  | Crow | -0.62278 | 1.075839 | -0.579 | 0.56267 |
|  | Finch | 0.300105 | 0.985515 | 0.305 | 0.76074 |
|  | Mouse | -1.57681 | 1.274309 | -1.237 | 0.21594 |
|  | White Noise | -1.81317 | 1.300463 | -1.394 | 0.16324 |
| Shelter Visits (Poisson) | Intercept | -0.84339 | 0.44038 | -1.915 | **0.0555** |
|  | Mass | -3.45183 | 2.30369 | -1.498 | 0.134 |
|  | Temp | -0.74589 | 0.44249 | -1.686 | 0.0919 |
|  | In Captivity | -0.0167 | 0.02577 | -0.648 | 0.517 |
|  | Injured Status | -0.45413 | 0.56727 | -0.801 | 0.4234 |
|  | Sex (M) | -1.2678 | 0.80955 | -1.566 | 0.1173 |
|  | Bat | -0.45951 | 0.58581 | -0.784 | 0.4328 |
|  | Crow | 0.47737 | 0.44455 | 1.074 | 0.2829 |
|  | Finch | -0.18883 | 0.56324 | -0.335 | 0.7374 |
|  | Mouse | -0.21896 | 0.59851 | -0.366 | 0.7145 |
|  | White Noise | -0.46668 | 0.58767 | -0.794 | 0.4271 |

| Table S4. Summary of results of pre-planned contrasts between treatments for total distance traveled, total time moving, and total time spent in shelter. P-values were adjusted using single-step method. Bold values indicate statistically significant effects (p<0.05). * indicates log transformed variables. | | | | | |
| --- | --- | --- | --- | --- | --- |
| Response Variable |  | Estimate | SE | z-value | p-value |
| Total Distance Traveled* | Control - Bat | -0.66731 | 0.24009 | -2.779 | **0.0481** |
|  | Control - Crow | -0.36632 | 0.23749 | -1.542 | 0.5841 |
|  | Control - Finch | -0.54364 | 0.25058 | -2.17 | 0.2125 |
|  | Control - Mouse | -0.74173 | 0.24255 | -3.058 | **0.021** |
|  | Control - White Noise | -0.69017 | 0.23253 | -2.968 | **0.0276** |
|  | White Noise - Bat | 0.02286 | 0.21941 | 0.104 | 1 |
|  | White Noise - Crow | 0.32385 | 0.22614 | 1.432 | 0.6587 |
|  | White Noise - Finch | 0.14653 | 0.23278 | 0.629 | 0.985 |
|  | White Noise- Mouse | -0.05156 | 0.22132 | -0.233 | 0.9999 |
|  | Crow - Finch | -0.17732 | 0.23888 | -0.742 | 0.969 |
|  | Bat - Mouse | -0.07442 | 0.23238 | -0.32 | 0.9994 |
| Total Time Moving | Control - Bat | -50.386 | 18.569 | -2.713 | 0.0577 |
|  | Control - Crow | -22.886 | 18.369 | -1.246 | 0.7757 |
|  | Control - Finch | -45.617 | 19.379 | -2.354 | 0.1425 |
|  | Control - Mouse | -53.523 | 18.759 | -2.853 | **0.039** |
|  | Control - White Noise | -56.438 | 17.986 | -3.138 | **0.0163** |
|  | White Noise - Bat | 6.052 | 16.969 | 0.357 | 0.9989 |
|  | White Noise - Crow | 33.552 | 17.49 | 1.918 | 0.3404 |
|  | White Noise - Finch | 10.821 | 18.001 | 0.601 | 0.9878 |
|  | White Noise- Mouse | 2.915 | 17.119 | 0.17 | 1 |
|  | Crow - Finch | -22.731 | 18.475 | -1.23 | 0.7847 |
|  | Bat - Mouse | -3.138 | 17.971 | -0.175 | 1 |
| Total Time in Shelter | Control - Bat | 127.54 | 32.1607 | 3.966 | **<0.001** |
|  | Control - Crow | 84.249 | 26.8867 | 3.133 | **0.0165** |
|  | Control - Finch | 83.2611 | 31.7053 | 2.626 | 0.0729 |
|  | Control - Mouse | 80.2918 | 31.6011 | 2.541 | 0.0909 |
|  | Control - White Noise | 90.1615 | 31.3224 | 2.878 | **0.0362** |
|  | White Noise - Bat | 37.3787 | 34.7505 | 1.076 | 0.866 |
|  | White Noise - Crow | -5.9125 | 30.6632 | -0.193 | 0.9999 |
|  | White Noise - Finch | -6.9004 | 34.7975 | -0.198 | 0.9999 |
|  | White Noise- Mouse | -9.8697 | 35.0179 | -0.282 | 0.9997 |
|  | Crow - Finch | -0.9879 | 29.2225 | -0.034 | 1 |
|  | Bat - Mouse | -47.2484 | 35.0795 | -1.347 | 0.7161 |

Table S5. Among-individual variances (V_A_), within-individual variances (V_W_), variation explained by fixed effects (V_F_), and adjusted repeatabilities (τ) for each behavioral response. * denotes log transformed values.

| Behavior | V_A_ (CI) | V_F_  (CI) | V_W_ (CI) | τ *(CI) |
| --- | --- | --- | --- | --- |
| Emergence Time* | 0.53 [0.113:1.073] | 0.299 [0.136:0.852] | 0.915 [0.636:1.21] | 0.367 [0.096:0.577] |
| Emergence | 7.157 [1.204:38.244] | 3.943 [1.521:32.408] | 4.5 [4.054:5.996] | 0.614 [0.172:0.874] |
| Total Distance* | 0.435 [0.142:0.832] | 0.171 [0.098:0.544] | 0.433 [0.296:0.583] | 0.501 [0.231:0.68] |
| Time Moving | 2557.532 [823.089:5085.007] | 824.644 [532.271:2917.6] | 2593.35 [1843.681:3538.679] | 0.497 [0.226:0.697] |
| Time Not Moving | 2796.073 [982.502:5580.455] | 1187.888 [597.867:3770.63] | 2968.15 [2076.868:4083.497] | 0.485 [0.188:0.673] |
| Freeze Response | 0 [NA:NA] | NA | NA | NA |
| Max Acceleration* | 0.1 [0.033:0.192] | 0.033 [0.017:0.125] | 0.098 [0.07:0.133] | 0.507 [0.223:0.684] |
| Time in Shelter | 1184.876 [0:3892.861] | 1841.779 [1405.426:4970.55] | 2625.198 [1179.26:4421.68] | 0.311 [0:0.707] |
| Returns to Shelter | 0.297 [0:0.703] | 0.623 [0.385:58.006] | 1.213 [0.946:1.807] | 0.197 [0:0.376] |
